# Identification of deleterious and regulatory genomic variations in known asthma loci

**DOI:** 10.1101/389031

**Authors:** Matthew D. C. Neville, Jihoon Choi, Jonathan Lieberman, Qing Ling Duan

**Author notes:** Corresponding Author: Qing Ling Duan, Address: Botterell Hall, Room 530 - 18 Stuart St, Kingston, ON K7L3N6.

## Abstract

**Background:** Candidate gene and genome-wide association studies have identified hundreds of asthma risk loci. The majority of associated variants, however, are not known to have any biological function and are believed to represent markers rather than true causative mutations. We hypothesized that many of these associated markers are in linkage disequilibrium (LD) with the elusive causative variants.

**Methods:** We compiled a comprehensive list of 447 asthma-associated variants previously reported in candidate gene and genome-wide association studies. Next, we identified all sequence variants located within the 304 unique genes using whole-genome sequencing data from the 1000 Genomes Project. Then, we calculated the LD between known asthma variants and the sequence variants within each gene. LD variants identified were then annotated to determine those that are potentially deleterious and/or functional (i.e. coding or regulatory effects on the encoded transcript or protein).

**Results:** We identified 10,048 variants in LD (r^2^ > 0.6) with known asthma variants. Annotations of these LD variants revealed that several have potentially deleterious effects including frameshift, alternate splice site, stop-lost, and missense. Moreover, 24 of the LD variants have been reported to regulate gene expression as expression quantitative trait loci (eQTLs).

**Conclusions:** This study is proof of concept that many of the genetic loci previously associated with complex diseases such as asthma are not causative but represent markers of disease, which are in LD with the elusive causative variants. We hereby report a number of potentially deleterious and regulatory variants that are in LD with the reported asthma loci. These reported LD variants could account for the original association signals with asthma and represent the true causative mutations at these loci.

## Background

Asthma is a chronic respiratory disease characterized by hyper-responsiveness of the bronchial muscles, inflammation, and reversible narrowing of the airways. It affects over 330 million individuals worldwide and is expected to increase to approximately 400 million by 2025 [1,2]. Asthma is known to be multifactorial and polygenic in nature with numerous genetic and environmental risk factors. Twin studies have estimated that genetic factors contribute approximately 55-95% of asthma risk, known as its heritability [3–5]. To date, hundreds of genetic variants have been correlated with asthma and asthma related traits such as lung function and bronchial hyper-responsiveness. A recent study of 31 of the most replicated loci for asthma, however, determined that these collectively account for only a small portion (2.6%) of asthma heritability [6]. Thus, our understanding of the genetic determinants of asthma risk remains limited and further investigations to identify the missing heritability are necessary to facilitate better prevention, diagnosis and care of this chronic disease.

Several factors may account of the missing heritability of asthma such as epistasis, gene-environment interactions, rare variants, and epigenomics (i.e. non-sequence variations in the genome) [7]. In addition, the genetic variants previously correlated with asthma may not represent the true causative mutation. Instead, it is widely accepted that the majority of loci identified from genome-wide association studies (GWAS) as well as candidate gene studies are not causal but are genetic markers that tag causal variants [8]. This is supported by the fact that the majority of variants associated with complex diseases such as asthma have no known impact on the resulting transcripts or proteins, with over 80% from GWAS falling outside of protein coding regions [9]. Discovery of the causal variants underlying a disease would reveal the true genetic effect sizes [8] and help to facilitate the development of more accurate clinical tests for diagnosis and treatment of asthma [10].

In this study, we hypothesize that the causal mutations at numerous asthma-associated loci are likely in linkage disequilibrium (LD) with the associated markers, which refers to the non-random association of alleles at different loci [11]. We test this through the analysis of existing whole-genome sequencing data from large populations to identify genetic variants that are more likely to be causal and explain for the missing heritability of asthma. First, we compiled a comprehensive list of asthma-associated variants from earlier GWAS and candidate gene studies. Next, we identified all variants within these asthma loci using sequence data from the 1000 Genomes Project (1000GP) [12] and calculated LD between the asthma variants and the sequence variants. Then, both asthma variants and the LD variants were annotated for their predicted effects on the resulting transcript or protein. We hereby report a list of potentially deleterious and regulatory variants within known asthma loci that are in LD with and could account for the reported association signals. These results may improve our understanding of the underlying mechanisms of asthma, which will ultimately lead to better prevention and more efficient therapies.

## Methods

### Selection of Asthma Loci

Previously associated SNPs from asthma genome-wide association studies (GWAS) were obtained from the National Human Genome Research Institute-European Bioinformatics Institute (NHGRI-EBI) GWAS Catalog (October 10^th^, 2017 release) [13]. This manually curated GWAS catalog lists SNP-trait associations with p-values < 1.0 x 10^-5^ in studies assaying at least 100,000 SNPs. We selected all SNPs linked to asthma and asthma severity traits: ‘Asthma’, ‘Adult asthma’, ‘Asthma (childhood onset)’, ‘Asthma (sex interaction)’, ‘Asthma and hay fever’, ‘Asthma or chronic obstructive pulmonary disease’, ‘Bronchial hyperresponsiveness in asthma’, and ‘Pulmonary function in asthmatics’.

A compilation of candidate genes were also selected from four previously published lists of asthma candidate gene studies [14–17]. A total of 148 genes that did not overlap with the asthma GWAS loci above were selected. We then proceeded to search for these candidate genes in PubMed using keywords including “asthma”, “polymorphism” or “snp”, and the gene name. All identified English language candidate gene studies reporting a genetic association for asthma or asthma severity published up to June 2016 were examined. Asthma-associated SNPs reported to a gene on our candidate gene list that were significant at the level defined by the authors of the study were included for further analysis.

### Genomic Coordinates and 1000 Genomes Project

Starting with a comprehensive list of single nucleotide polymorphisms previously associated with asthma and related phenotypes from GWAS and candidate gene studies, we removed sex chromosome genes, pseudogenes and uncharacterized loci unknown to the RefSeqGene database [18]. Next, the University of Santa Cruz (UCSC) Genome Browser [19] was used to identify the genomic coordinates of each gene using GRCh37/hg19 assembly of the human reference genome. The coordinates of each gene were extended by 5000 bp both 5’ and 3’ of the transcription start sites (TSS) and transcription end sites (TES) of each gene to include potential regulatory regions. These genomic coordinates were then applied to extract a complete list of sequence variations from the 1000GP Phase 3 whole genome sequencing data [12].

### Linkage Disequilibrium Analysis

At each locus, we calculated the LD between the asthma-associated SNP (reported by GWAS and candidate gene studies) and the newly identified variants from the 1000GP using Plink, version 1.9 [20]. A distance window of 1Mb was used to determine LD, which excludes variants greater than 1Mb apart. The r^2^ threshold for LD was set to ≥ 0.6.

### Functional Annotation

The genomic variant annotations and functional effect prediction toolbox known as SnpEff (version 4.3q) [21] was used with the GRCh37.75 assembly to predict the effects of sequence variants identified from the 1000GP. To validate these annotations, we also used Ensembl’s Variant Effect Predictor (VEP), release 90, [22] and only concordant annotations between the two prediction tools were reported. For those variants with multiple annotations from each annotation tool (e.g. variants in regions impacted by multiple transcripts), only the most severe effect as ranked by SnpEff was selected. VEP was also used to access the pathogenicity predictions of missense mutations from the scoring tools Sorting Intolerant From Tolerant (SIFT) [23] and Polyphen-2 [24]. The Database of Genomic Structural Variations (dbVar) [25] was used to supplement annotations for structural variants.

RegulomeDB (version 1.1) [26] was used to evaluate the potential of variants to impact gene expression. RegulomeDB scores variants for their predicted functional impact based on high-throughput data from non-coding and intergenic regions of the human genome. It uses experimental data sets from the Encyclopedia of DNA Elements (ENCODE) project [27], public datasets from Gene Expression Omnibus (GEO) [28], and curated published literature.

## Results

### Asthma Loci

We identified a total of 447 asthma-associated SNPs from earlier GWAS and candidate gene studies. Of these, 225 SNPs were found from asthma GWAS via the NHGRI-EBI GWAS Catalog [13] and mapped to 224 unique genes (**Additional file 1: Table S1**). In addition, we identified loci reported by candidate gene studies, of which 222 SNPs did not overlap with GWAS loci and mapped to 80 unique genes (**Additional file 1: Table S2**). Functional annotation of the 447 asthma variants revealed that only 12% are predicted to cause protein coding changes (**Additional file 1: Table S3**), while most are intronic (30%), intergenic (15%), or upstream/downstream (32%) gene variants (**Fig. 1**).

**Fig. 1.**
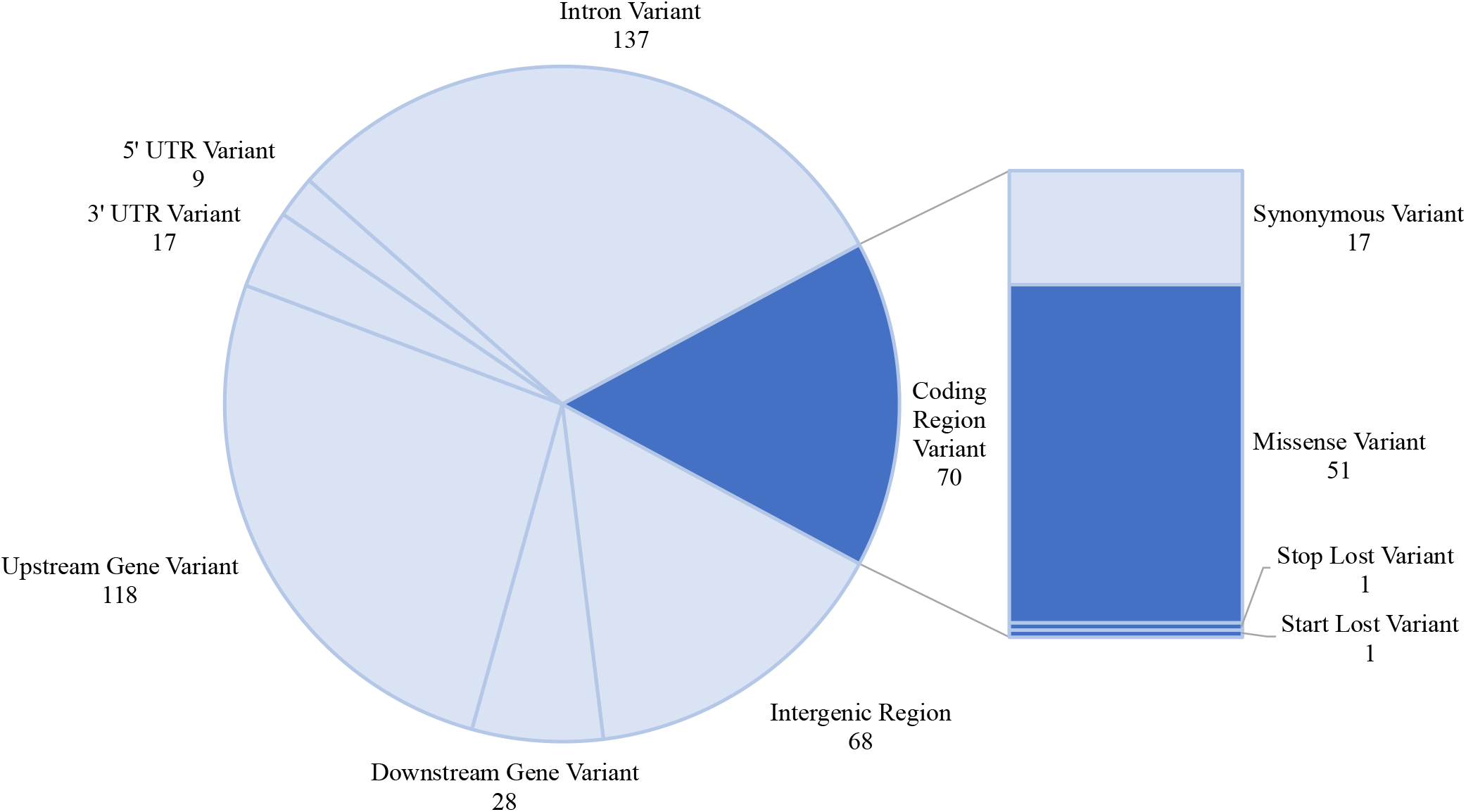
Number of asthma-associated variants by annotation. Genomic variants were previously reported by genome-wide association or candidate gene studies of asthma and asthma related traits. Annotations derived from SnpEff indicate that approximately 12% of the known asthma variants code for a missense or non-sense variant, whereas the majority are non-coding or have no known function.

### Linkage Disequilibrium Variants

Using whole-genome sequencing data within the genomic coordinates of the 304 unique asthma genes from GWAS and candidate gene studies, we identified potentially functional variants that are in LD with asthma-associated variants. Specifically, we extracted a total of 1,382,700 variants from these 304 loci using Phase 3 sequence data of the 1000GP. We then assessed LD between the asthma variants and the sequence variants from the 1000GP using data from each of the five continental ancestry groups (African, American, East Asian, European, and South Asian) to account for the discordant nature of LD among ethnic groups [29,30]. All variants in LD (r^2^ ≥ 0.6) in one or more of the five ancestry groups were included for further analysis. This resulted in 10,048 variants forming 14,826 instances of LD (r^2^ ≥ 0.6) with 342 asthma-associated variants (**Additional file 1: Table S4**). These 10,048 variants from the 1000GP in LD with one or more asthma variants are hereafter referred to as the ‘LD variants’ and the workflow to generate and annotate them is summarized in **Fig. 2**. Among the LD variants, 9,073 are SNPs, 966 are insertions or deletions, and 9 are structural variants (**Additional file 1: Table S5**).

**Fig. 2.**
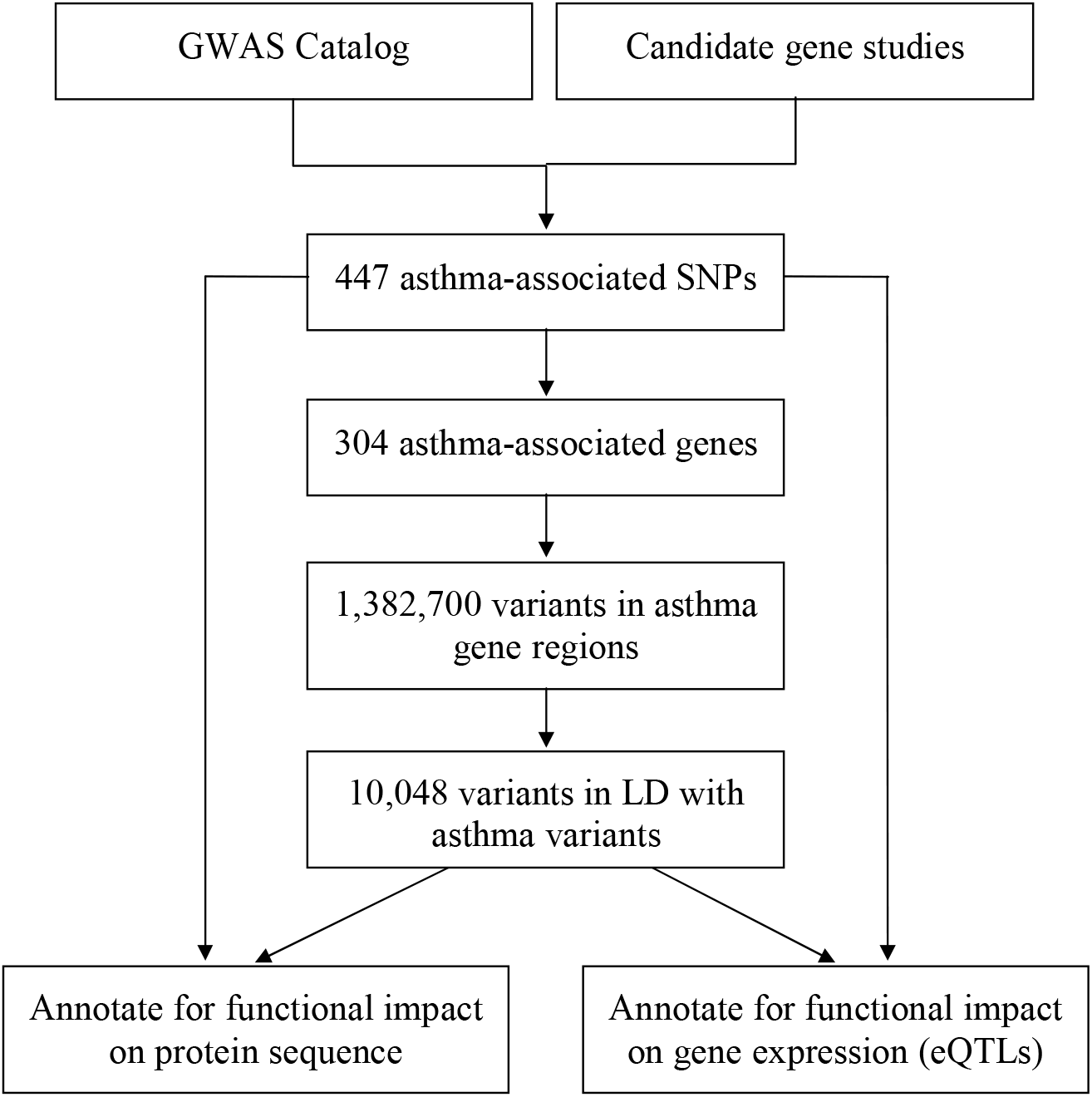
Workflow for discovery of potentially causal variants in known asthma loci. First, we compiled and annotated asthma-associated SNPs from previous GWAS and candidate gene studies of asthma. We then determined the genomic coordinates of the 304 genes and extracted all sequence variations within these loci using data from the 1000 Genomes Project (1000GP). Finally, we identified and annotated those variants in linkage disequilibrium (LD) with known asthma loci to determine which may contribute to changes in protein sequence or regulate gene expression (i.e. expression quantitative trait loci or eQTL).

### Identification of deleterious and regulatory variants

Many of the LD variants are predicted to have functional consequences on resulting protein structure or to impact regulation of gene expression. **Table 1** reports 6 LD variants that are predicted to result in frameshift, stop lost, or splice site mutations by both genomic annotation tools SnpEff and VEP. Two deletions (rs199503730 and rs67841474) are annotated as frameshift variants, both impacting the final coding exon of major histocompatibility complex (MHC) Class I gene *MICA*. The deletion variant rs146576636 in the final coding exon of *ADAM33* is also annotated as a frameshift variant. Two SNPs (rs8084 and rs11078928) are annotated as splice acceptor variants in *HLA-DRA* and *GSDMB* respectively. Finally, the SNP rs15895 is annotated as a stop lost variant in *OAS2*. These variants are in LD with previously reported asthma variants that are less likely to be functional, such as intron variants, upstream/downstream variants, or synonymous variants.

**Table 1.**
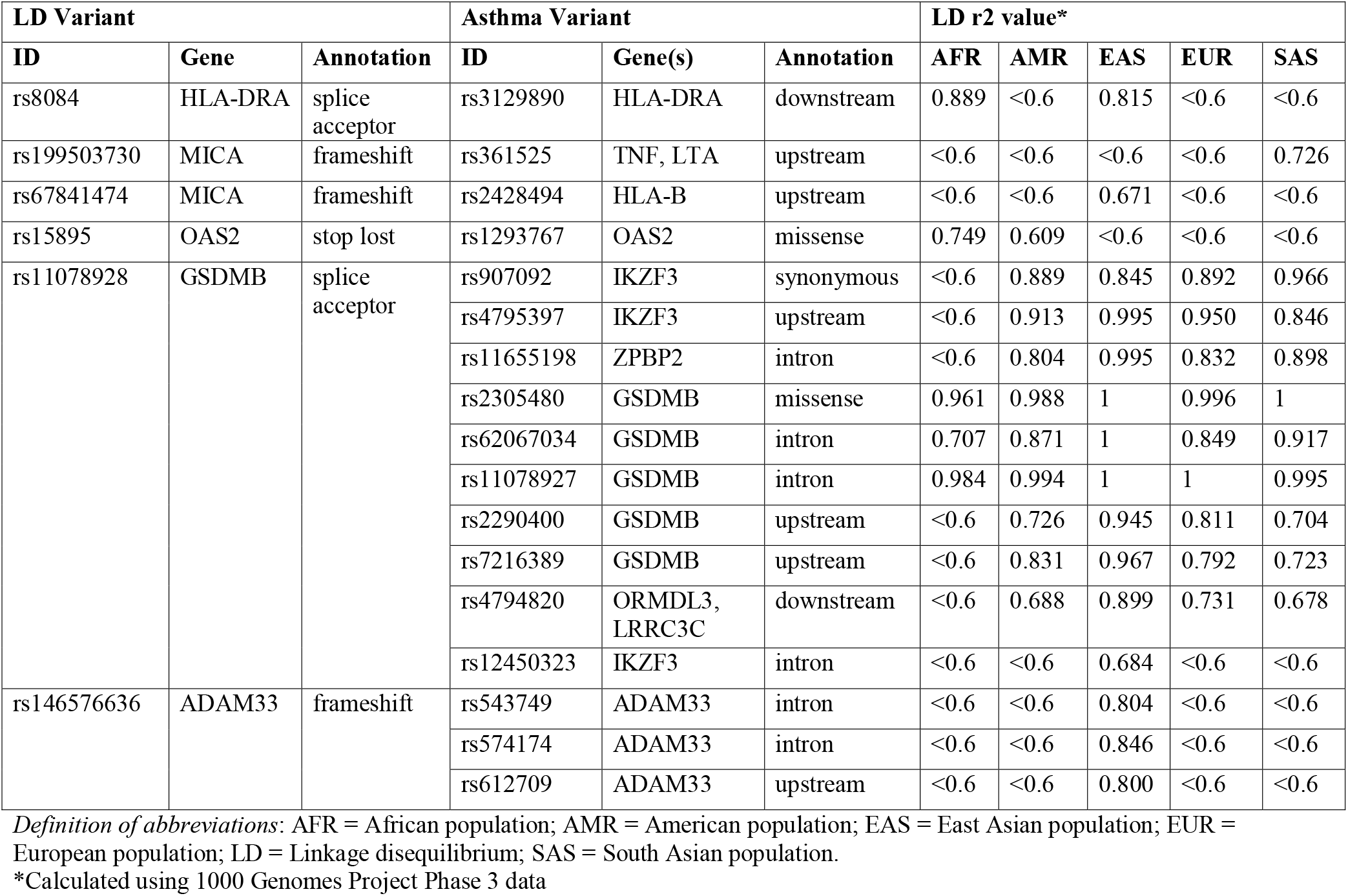
Deleterious coding variants in LD with asthma-associated SNPs.

In addition to these potentially deleterious coding variants, we identified 91 LD variants that are predicted to be missense variants by both SnpEff and VEP. Of these, 34 are classified as deleterious and/or probably damaging by the pathogenicity prediction tools SIFT and Polyphen-2 (**Table 2**). The remaining 57 missense LD variants were not identified as deleterious or probably damaging using these software (**Additional file 1: Table S6**).

**Table 2.** Missense variants in LD with asthma-associated variants

| LD Variants |  |  | Asthma Variants |
| --- | --- | --- | --- |
| ID | Gene | Annotation | ID(s) |
| rs115676129 | ABI3BP | missense<sup>*,†</sup> | rs9823506 |
| rs2275254 | CHIA | missense<sup>*,†</sup> | rs2282290 |
| rs2305479 | GSDMB | missense<sup>*,†</sup> | rs2305480, rs62067034, rs11078927, rs7216389, rs907092, rs4795397, rs11655198, rs2290400, rs12450323, rs4794820 |
| rs1129808 | HLA-DQA1 | missense<sup>*,†</sup> | rs3104367, rs9272346, rs9273373 |
| rs1142332 | HLA-DQA1 | missense<sup>*,†</sup> | rs3104367, rs9272346, rs9273373 |
| rs1049057 | HLA-DQB1 | missense<sup>*,†</sup> | rs3104367, rs9272346, rs9273373 |
| rs1140319 | HLA-DQB1 | missense<sup>*,†</sup> | rs3104367, rs9272346, rs9273373 |
| rs73022563 | PLEKHG4B | missense<sup>*,†</sup> | rs62344088 |
| rs6482626 | PTCHD3 | missense<sup>*,†</sup> | rs660498 |
| rs2305089 | TBXT | missense<sup>*,†</sup> | rs6456042 |
| rs41273547 | ABI3BP | missense<sup>*</sup> | rs9823506 |
| rs148714608 | CLSTN2 | missense<sup>*</sup> | rs77960860 |
| rs1042308 | HLA-DPA1 | missense<sup>*</sup> | rs987870 |
| rs1071630 | HLA-DQA1 | missense<sup>*</sup> | rs3104367, rs9272346, rs9273373 |
| rs1129740 | HLA-DQA1 | missense<sup>*</sup> | rs3104367, rs9272346, rs9273373 |
| rs1142328 | HLA-DQA1 | missense<sup>*</sup> | rs3104367, rs9272346, rs9273373 |
| rs1142331 | HLA-DQA1 | missense<sup>*</sup> | rs3104367, rs9272346, rs9273373 |
| rs3208105 | HLA-DQA1 | missense<sup>*</sup> | rs3104367, rs9272346, rs9273373 |
| rs1049163 | HLA-DQB1 | missense<sup>*</sup> | rs9273373 |
| rs1130398 | HLA-DQB1 | missense<sup>*</sup> | rs3104367, rs9272346, rs9273373 |
| rs1140318 | HLA-DQB1 | missense<sup>*</sup> | rs3104367, rs9272346, rs9273373 |
| rs1140320 | HLA-DQB1 | missense<sup>*</sup> | rs3104367, rs9272346, rs9273373 |
| rs41558312 | MICA | missense<sup>*</sup> | rs361525 |
| rs1801280 | NAT2 | missense<sup>*</sup> | rs1799929, rs1208 |
| rs2363468 | PIKFYVE | missense<sup>*</sup> | rs4673397 |
| rs13436090 | PLEKHG4B | missense<sup>*</sup> | rs62344088 |
| rs3777134 | SPINK5 | missense<sup>*</sup> | rs2303064 |
| rs682632 | TEK | missense<sup>*</sup> | rs72721168 |
| rs1140404 | HLA-B | missense<sup>†</sup> | rs1800629 |
| rs41543121 | HLA-B | missense<sup>†</sup> | rs1800629 |
| rs41556417 | HLA-B | missense<sup>†</sup> | rs361525 |
| rs4193 | HLA-DQA1 | missense<sup>†</sup> | rs3104367, rs9272346, rs9273373 |
| rs2233953 | PSORS1C2 | missense<sup>†</sup> | rs1800629 |
| rs7686508 | SEC31A | missense<sup>†</sup> | rs10022260 |

*Definition of abbreviations:* LD = Linkage disequilibrium
\*Predicted to be deleterious by Sorting Intolerant From Tolerant (SIFT)
†Predicted to be probably damaging by Polyphen-2

Finally, we identified LD variants that have been associated with gene expression (i.e. expression quantitative trait loci (eQTL)) using RegulomeDB. We limited our investigation to eQTLs located both at a transcription factor binding site and a DNase peak, two factors characteristic of causal eQTLs [31]. In total, we determined that 24 of the LD variants (**Table 3**) and 7 of the asthma-associated variants (**Additional file 1: Table S7**) are known eQTLs.

**Table 3.**
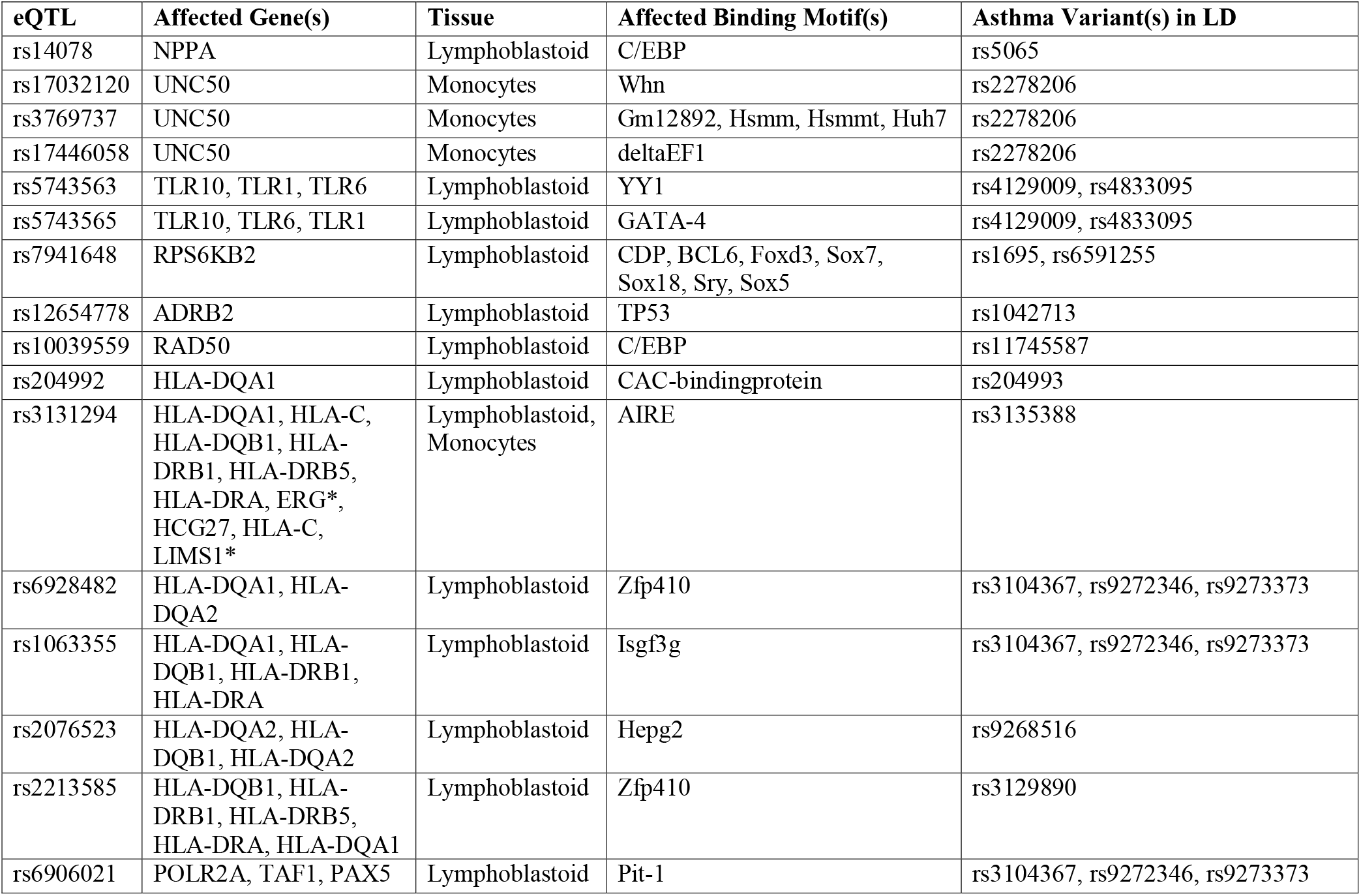
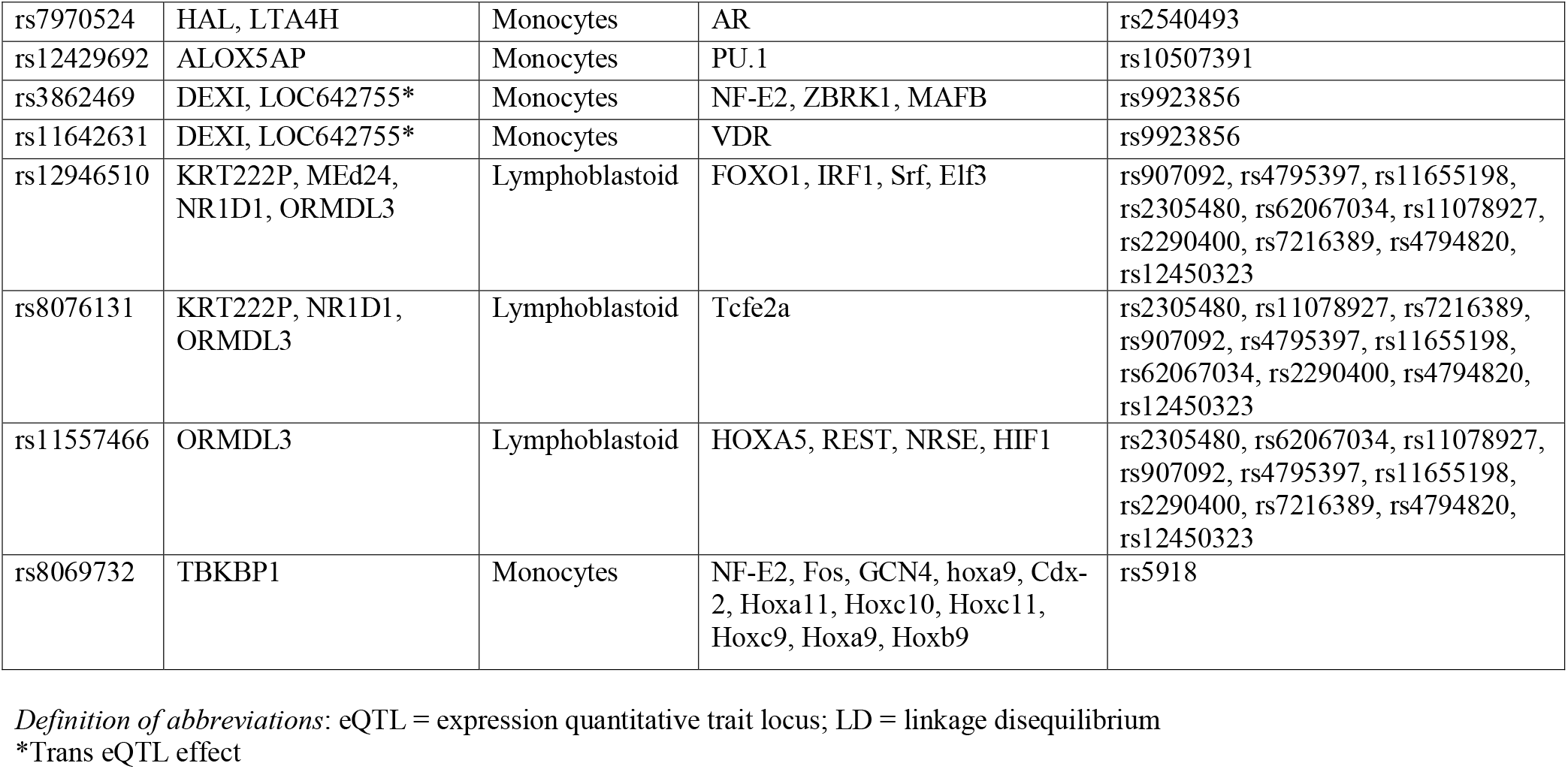
Regulatory variants (eQTLs) in LD with asthma-associated variants

## Discussion

In this study, we compiled a comprehensive list of asthma-associated variants from earlier candidate gene studies and GWAS in order to conduct a systematic search for causal variants that likely contribute to asthma heritability. Whole genome sequencing data from the 1000GP and functional annotation software tools were used to identify potentially deleterious variants in LD with the asthma-associated variants that could account for the original association signals at these loci. We identified variants annotated to be frameshift, splice site, stop-lost, missense, or eQTLs that are in LD with known asthma variants. The majority of these functional variants were in LD with asthma variants that are non-coding and have no known effects on gene transcription. In summary, we have identified numerous functional variants that could be the elusive causal variants within known asthma-associated loci and improve our understanding of the underlying mechanisms of asthma.

Several of the potentially causal variants identified in this study are found in the MHC class II locus, within the human leukocyte antigen (HLA) super-locus of the 6p21 chromosomal region. MHC class II genes code for proteins that play a central role in the immune system by presenting peptides to CD4+ T cells [32]. The MHC class II locus has been repeatedly associated to asthma and other immune related diseases, but finding causal variants at the locus has proven to be difficult due to its highly polymorphic nature and strong linkage disequilibrium in the region [33,34]. We compiled 10 asthma risk variants (rs9268516, rs9272346, rs9273349, rs9273373, rs9275698, rs9500927, rs987870, rs3104367, rs3129890, rs7775228) found at this locus from seven GWAS [35–41]. Only two of these asthma-associated variants (rs9272346 and rs7775228) have been reported to regulate gene expression (eQTLs), whereas the remainder are not known to have any functional impact on the resultant transcripts or proteins. The present study identified numerous LD variants within the same loci as these asthma variants including a splice-acceptor variant in the *HLA-DRA* gene (rs8084); missense variants in *HLA-DQA1, HLA-DQB1, HLA-DRB, HLA-DRB5*, and *HLA-DPA1*; and several eQTLs (rs204992, rs3131294, rs6928482, rs1063355, rs2076523, rs2213585) that are linked to the expression of one or more MHC class II genes. Taken together these LD variants, which are more likely to have functional or even detrimental consequences, could account for the consistent associations between the MHC class II locus and asthma outcomes.

A number of the variants identified in this study are located within the 17q12-21 chromosomal region, which is the most reproducible asthma locus [42,43]. This genomic region consists of a number of genes, including *ORMDL3, GSDMB, IKZF3, ZPBP2*, and *GSDMA* that are in LD and may either be individually or jointly responsible for the asthma association [42]. The current study compiled 13 asthma-associated variants from the 17q12-21 region (rs3894194, rs4795397, rs62067034, rs7212938, rs7216389, rs11078927, rs2290400, rs2305480, rs4794820, rs6503525, rs11655198, rs12450323, rs907092) that were identified across 11 independent GWAS [37–39,43–50]. Most of these variants are non-coding, save three missense mutations: rs3894194 and rs7212938 from *GSDMA* and rs2305480 from *GSDMB*. This study identified four additional coding variants from *GSDMA* (rs56030650), *GSDMB* (rs2305479, rs11078928) and *ZPBP2* (rs11557467) and three eQTLs for *ORMDL3* (rs12946510, rs11557466, rs8076131), which are in LD with asthma variants. Notably the LD variant rs11078928 has been functionally characterized as a splice variant that removes an exon, influences transcription levels, and abolishes the biochemical activity of *GSDMB* [51,52]. This variant has been previously discussed as a possible causal variant in asthma [42,53] and its LD with multiple asthma-associated variants supports such a hypothesis. The LD variants identified in the 17q12-21 locus demonstrate the possibility that these could represent the causal variants underlying asthma risk.

While we report potentially causal variants in known asthma loci, our study also has some important limitations. For example, given that LD calculations are dependent on allele frequencies and that the majority of known asthma variants are common (minor allele frequency or MAF > 0.05), our study is biased toward identifying LD variants with higher MAF [54]. Additionally, our list of asthma-associated variants from earlier GWAS and candidate gene studies may contain some false positive results. Further studies are needed to validate these associations in additional asthma populations. Finally, another limitation of our study is that we relied on annotation data from prediction tools such as SnpEff and VEP, which are limited and do not replace experimental evidence. Further studies are needed to identify and ultimately validate the true function, if any, of the LD variants identified in this study to be coding or regulatory. These additional studies include genotyping of these potentially functional variants (identified in LD with the associated variants) in asthma populations and testing them directly for correlation with asthma outcomes. Moreover, functional validation experiments are needed to confirm the biological impact of these variants on the resultant RNA transcripts or proteins, which depends on the predicted impact of the variants identified. For example, variants of high impact (**Table 1**) include frameshift, splice variants and premature stop codons, could be validated through direct sequencing of transcripts and mass spectrometry to detect truncated and mis-folded proteins.

## Conclusions

The current study is proof of concept that previously correlated loci for asthma may tag causal variants that are in LD, which can be identified using direct sequencing data. While genetic studies of asthma to date have successfully identified hundreds of asthma loci, our understanding of the underlying mechanisms of asthma remains limited. In addition, these loci do not account for all of the asthma heritability and have limited clinical applications due to the fact that the majority of associated variants do not represent the true causative loci. Identification of these causal variants underlying the genetic associations will be instrumental in improving our understanding of the underlying mechanisms of asthma. In addition, knowledge of the casual variants will ultimately help to facilitate the development of more accurate clinical tests for determining risk and treatment options for asthma.

## Abbreviations

1000GP: : 1000 Genomes Project
eQTL: : Expression quantitative trait loci
GWAS: : Genome-wide association study
HLA: : Human leukocyte antigen
LD: : Linkage disequilibrium
MHC: : Major histocompatibility complex
NHGRI-EBI: : National Human Genome Research Institute-European Bioinformatics Institute
SIFT: : Sorting Intolerant From Tolerant
SNP: : Single nucleotide polymorphism
VEP: : Variant Effect Predictor

## Declarations

### Ethics approval and consent to participate

Not applicable.

### Consent for publication

Not applicable.

### Availability of data and material

The 1000 Genomes Project Phase 3 dataset used for the current analyses is publicly available from The International Genome Sample Resource (http://www.internationalgenome.org/).

### Authors' contributions

Conception and design: QLD. Data analysis: MN, JL, JC. Wrote manuscript: MN, JC, QLD. Critical revision of the manuscript: all authors. All authors read and approved the final manuscript.

## Acknowledgements

The authors thank the Centre for Advanced Computing (CAC) at Queen's University in Kingston, Ontario for computational resources and support. The CAC is funded by: the Canada Foundation for Innovation, the Government of Ontario, and Queen's University.

## Supplementary Material

**Additional file 1: Table S1.** Asthma-associated SNPs from genome-wide association studies. **Table S2.** Asthma-associated SNPs from candidate gene studies. **Table S3.** Asthma-associated variants with protein coding annotation. **Table S4.** Ethnic specific linkage disequilibrium between asthma variants and 1000GP variants. **Table S5.** Annotation of all variants in linkage disequilibrium with asthma variants. **Table S6.** Missense variants not predicted to be deleterious in linkage disequilibrium with asthma-associated variants. **Table S7.** eQTLs within asthma-associated variants.

## References

1 Vos T, Flaxman AD, Naghavi M, et al. Years lived with disability (YLDs) for 1160 sequelae of 289 diseases and injuries 1990–2010: a systematic analysis for the Global Burden of Disease Study 2010. Lancet 2012;380:2163–96. doi:10.1016/S0140-6736(12)61729-2

2 Masoli M, Fabian D, Holt S, et al. The global burden of asthma: executive summary of the GINA Dissemination Committee Report. Allergy 2004;59:469–78. doi:10.1111/j.1398-9995.2004.00526.x

3 Duffy DL, Martin NG, Battistutta D, et al. Genetics of Asthma and Hay Fever in Australian Twins. Am Rev Respir Dis 1990;142:1351–8. doi:10.1164/ajrccm/142.6_Pt_1.1351

4 Nieminen MM, Kaprio J, Koskenvuo M. A population-based study of bronchial asthma in adult twin pairs. Chest 1991;100:70–5. http://www.ncbi.nlm.nih.gov/pubmed/2060393 (accessed 21 Dec 2017).

5 Thomsen SF, Van Der Sluis S, Kyvik KO, et al. Estimates of asthma heritability in a large twin sample Clinical & Experimental Allergy. Clin Exp Allergy 2010;1054–61. doi:10.1111/j.1365-2222.2010.03525.x

6 Vicente CT, Revez JA, Ferreira MAR. Lessons from ten years of genome-wide association studies of asthma. Clin Transl Immunol 2017;6:e165. doi:10.1038/cti.2017.54

7 Ober C, Yao T-C. The genetics of asthma and allergic disease: a 21st century perspective. Immunol Rev 2011;242:10–30. doi:10.1111/j.1600-065X.2011.01029.x

8 Manolio TA, Collins FS, Cox NJ, et al. Finding the missing heritability of complex diseases. Nature 2009;461:747–53. doi:10.1038/nature08494

9 Hindorff LA, Sethupathy P, Junkins HA, et al. Potential etiologic and functional implications of genome-wide association loci for human diseases and traits. Proc Natl Acad Sci U S A 2009;106:9362–7. doi:10.1073/pnas.0903103106

10 MacArthur DG, Manolio TA, Dimmock DP, et al. Guidelines for investigating causality of sequence variants in human disease. Nature 2014;508:469–76. doi:10.1038/nature13127

11 Pritchard JK, Przeworski M. Linkage disequilibrium in humans: models and data. Am J Hum Genet 2001;69:1–14. doi:10.1086/321275

12 Auton A, Abecasis GR, Altshuler DM, et al. A global reference for human genetic variation. Nature 2015;526:68–74. doi:10.1038/nature15393

13 MacArthur J, Bowler E, Cerezo M, et al. The new NHGRI-EBI Catalog of published genome-wide association studies (GWAS Catalog). Nucleic Acids Res 2017;45:D896–901. doi:10.1093/nar/gkw1133

14 Melén E, Kho AT, Sharma S, et al. Expression analysis of asthma candidate genes during human and murine lung development. Respir Res 2011;12:86. doi:10.1186/1465-9921-12-86

15 Sharma A, Menche J, Huang CC, et al. A disease module in the interactome explains disease heterogeneity, drug response and captures novel pathways and genes in asthma. Hum Mol Genet 2015;24:3005–20. doi:10.1093/hmg/ddv001

16 Vercelli D. Discovering susceptibility genes for asthma and allergy. Nat Rev Immunol 2008;8:169–82. doi:10.1038/nri2257

17 Weiss ST, Raby BA, Rogers A. Asthma genetics and genomics 2009. Curr Opin Genet Dev 2009;19:279–82. doi:10.1016/j.gde.2009.05.001

18 Pruitt KD, Brown GR, Hiatt SM, et al. RefSeq: an update on mammalian reference sequences. Nucleic Acids Res 2014;42:D756–63. doi:10.1093/nar/gkt1114

19 Karolchik D, Hinrichs AS, Furey TS, et al. The UCSC Table Browser data retrieval tool. Nucleic Acids Res 2004;32:493D–496. doi:10.1093/nar/gkh103

20 Chang CC, Chow CC, Tellier LC, et al. Second-generation PLINK: rising to the challenge of larger and richer datasets. Gigascience 2015;4:7. doi:10.1186/s13742-015-0047-8

21 Cingolani P, Platts A, Wang LL, et al. A program for annotating and predicting the effects of single nucleotide polymorphisms, SnpEff: SNPs in the genome of Drosophila melanogaster strain w1118; iso-2; iso-3. Fly (Austin) 2012;6:80–92. doi:10.4161/fly.19695

22 McLaren W, Gil L, Hunt SE, et al. The Ensembl Variant Effect Predictor. Genome Biol 2016;17:122. doi:10.1186/s13059-016-0974-4

23 Kumar P, Henikoff S, Ng PC. Predicting the effects of coding non-synonymous variants on protein function using the SIFT algorithm. Nat Protoc 2009;4:1073–81. doi:10.1038/nprot.2009.86

24 Adzhubei IA, Schmidt S, Peshkin L, et al. A method and server for predicting damaging missense mutations. Nat Methods 2010;7:248–9. doi:10.1038/nmeth0410-248

25 Lappalainen I, Lopez J, Skipper L, et al. DbVar and DGVa: public archives for genomic structural variation. Nucleic Acids Res 2013;41:D936–41. doi:10.1093/nar/gks1213

26 Boyle AP, Hong EL, Hariharan M, et al. Annotation of functional variation in personal genomes using RegulomeDB. Genome Res 2012;22:1790–7. doi:10.1101/gr.137323.112

27 ENCODE Project Consortium TEP. An integrated encyclopedia of DNA elements in the human genome. Nature 2012;489:57–74. doi:10.1038/nature11247

28 Barrett T, Wilhite SE, Ledoux P, et al. NCBI GEO: archive for functional genomics data sets—update. Nucleic Acids Res 2012;41:D991–5. doi:10.1093/nar/gks1193

29 Charles BA, Shriner D, Rotimi CN. Accounting for Linkage Disequilibrium in Association Analysis of Diverse Populations. Genet Epidemiol 2014;38:265–73. doi:10.1002/gepi.21788

30 Ong RTH, Teo YY. varLD: a program for quantifying variation in linkage disequilibrium patterns between populations. Bioinformatics 2010;26:1269–70. doi:10.1093/bioinformatics/btq125

31 Albert FW, Kruglyak L. The role of regulatory variation in complex traits and disease. Nat Rev Genet 2015;16:197–212. doi:10.1038/nrg3891

32 Watts C. The exogenous pathway for antigen presentation on major histocompatibility complex class II and CD1 molecules. Nat Immunol 2004;5:685–92. doi:10.1038/ni1088

33 Kontakioti E, Domvri K, Papakosta D, et al. HLA and asthma phenotypes/endotypes: A review. Hum Immunol 2014;75:930–9. doi:10.1016/J.HUMIMM.2014.06.022

34 Handunnetthi L, Ramagopalan S V, Ebers GC, et al. Regulation of major histocompatibility complex class II gene expression, genetic variation and disease. Genes Immun 2010;11:99–112. doi:10.1038/gene.2009.83

35 Ramasamy A, Kuokkanen M, Vedantam S, et al. Genome-wide association studies of asthma in population-based cohorts confirm known and suggested loci and identify an additional association near HLA. PLoS One 2012;7:e44008. doi:10.1371/journal.pone.0044008

36 Lasky-Su J, Himes BE, Raby BA, et al. HLA-DQ strikes again: genome-wide association study further confirms HLA-DQ in the diagnosis of asthma among adults. Clin Exp Allergy 2012;42:1724–33. doi:10.1111/cea.12000

37 Moffatt MF, Gut IG, Demenais F, et al. A large-scale, consortium-based genomewide association study of asthma. N Engl J Med 2010;363:1211–21. doi:10.1056/NEJMoa0906312

38 Pickrell JK, Berisa T, Liu JZ, et al. Detection and interpretation of shared genetic influences on 42 human traits. Nat Genet 2016;48:709–17. doi:10.1038/ng.3570

39 Ferreira MAR, Matheson MC, Tang CS, et al. Genome-wide association analysis identifies 11 risk variants associated with the asthma with hay fever phenotype. J Allergy Clin Immunol 2014;133:1564–71. doi:10.1016/j.jaci.2013.10.030

40 Noguchi E, Sakamoto H, Hirota T, et al. Genome-wide association study identifies HLA-DP as a susceptibility gene for pediatric asthma in Asian populations. PLoS Genet 2011;7:e1002170. doi:10.1371/journal.pgen.1002170

41 Hirota T, Takahashi A, Kubo M, et al. Genome-wide association study identifies three new susceptibility loci for adult asthma in the Japanese population. Nat Genet 2011;43:893–6. doi:10.1038/ng.887

42 Stein MM, Thompson EE, Schoettler N, et al. A decade of research on the 17q12-21 asthma locus: Piecing together the puzzle. J Allergy Clin Immunol Published Online First: 4 January 2018. doi:10.1016/J.JACI.2017.12.974

43 Torgerson DG, Ampleford EJ, Chiu GY, et al. Meta-analysis of genome-wide association studies of asthma in ethnically diverse North American populations. Nat Genet 2011;43:887–92. http://www.ncbi.nlm.nih.gov/pubmed/21804549 (accessed 28 Nov 2017).

44 Wan YI, Shrine NRG, Soler Artigas M, et al. Genome-wide association study to identify genetic determinants of severe asthma. Thorax 2012;67:762–8. doi:10.1136/thoraxjnl-2011-201262

45 Moffatt MF, Kabesch M, Liang L, et al. Genetic variants regulating ORMDL3 expression contribute to the risk of childhood asthma. Nature 2007;448:470–3. doi:10.1038/nature06014

46 Yan Q, Brehm J, Pino-Yanes M, et al. A meta-analysis of genome-wide association studies of asthma in Puerto Ricans. Eur Respir J 2017;49:1601505. doi:10.1183/13993003.01505-2016

47 Nieuwenhuis MA, Siedlinski M, van den Berge M, et al. Combining genomewide association study and lung eQTL analysis provides evidence for novel genes associated with asthma. Allergy 2016;71:1712–20. doi:10.1111/all.12990

48 Almoguera B, Vazquez L, Mentch F, et al. Identification of Four Novel Loci in Asthma in European American and African American Populations. Am J Respir Crit Care Med 2017;195:456–63. doi:10.1164/rccm.201604-0861OC

49 Bønnelykke K, Sleiman P, Nielsen K, et al. A genome-wide association study identifies CDHR3 as a susceptibility locus for early childhood asthma with severe exacerbations. Nat Genet 2013;46:51–5. doi:10.1038/ng.2830

50 Ferreira MAR, McRae AF, Medland SE, et al. Association between ORMDL3, IL1RL1 and a deletion on chromosome 17q21 with asthma risk in Australia. Eur J Hum Genet 2011;19:458–64. doi:10.1038/ejhg.2010.191

51 Morrison FS, Locke JM, Wood AR, et al. The splice site variant rs11078928 may be associated with a genotype-dependent alteration in expression of GSDMB transcripts. BMC Genomics 2013;14:627. doi:10.1186/1471-2164-14-627

52 Panganiban RA, Sun M, Dahlin A, et al. A functional splice variant associated with decreased asthma risk abolishes the ability of gasdermin B to induce epithelial cell pyroptosis. J Allergy Clin Immunol Published Online First: 9 January 2018. doi:10.1016/j.jaci.2017.11.040

53 Igartua C, Myers RA, Mathias RA, et al. Ethnic-specific associations of rare and low-frequency DNA sequence variants with asthma. Nat Commun 2015;6:5965. doi:10.1038/ncomms6965

54 VanLiere JM, Rosenberg NA. Mathematical properties of the r2 measure of linkage disequilibrium. Theor Popul Biol 2008;74:130–7. doi:10.1016/j.tpb.2008.05.006

